# PyIOmica: Longitudinal Omics Analysis and Classification

**DOI:** 10.1101/708941

**Authors:** Sergii Domanskyi, Carlo Piermarocchi, George I. Mias

## Abstract

**Summary:** PyIOmica is an open-source Python package focusing on integrating longitudinal multiple omics datasets, characterizing, and classifying temporal trends. The package includes multiple bioinformatics tools including data normalization, annotation, classification, visualization, and enrichment analysis for gene ontology terms and pathways. Additionally, the package includes an implementation of visibility graphs to visualize time series as networks.

**Availability and implementation:** PyIOmica is implemented as a Python package (pyiomica), available for download and installation through the Python Package Index (PyPI) (https://pypi.python.org/pypi/pyiomica), and can be deployed using the Python import function following installation. PyIOmica has been tested on Mac OS X, Unix/Linux and Microsoft Windows. The application is distributed under an MIT license. Source code for each release is also available for download on Zenodo (https://doi.org/10.5281/zenodo.3342612).

## 1 Introduction

As sequencing costs continue to drop, systems biology based on large omics datasets is rapidly expanding its scope. In particular, time series obtained from multi-omics datasets are becoming more and more affordable. The analysis of time series can have broad implications for precision medicine applications, since longitudinal data capture the dynamically-changing collective microscopic behavior of molecular components in the body, reflecting the physiological state of a patient. There are many bioinformatics tools aiming at multimodal omics data integration (Pinu *et al*., 2019). Specifically, Bioconductor (Gentleman *et al*., 2004), Galaxy(Afgan *et al*., 2018), GenePattern(Reich *et al*., 2006), Biopython (Cock *et al*., 2009), Pathomx (Fitzpatrick *et al*., 2014), SECIMTools (Kirpich *et al*., 2018), and more. While multiple coding paradigms are used in bioinformatics, R and Python are essentially the lingua francas for data science analysis, where the open-source appeal and growing online community support are particularly helpful in developing a dedicated user base.

Here we introduce PyIOmica, an open source Python package, for analyzing longitudinal omics datasets, which includes multiple tools for processing of multi-modal mapped data, characterizing time series in terms of periodograms and autocorrelations, classifying temporal behavior, visualizing visibility graphs, and testing data for gene ontology and pathway enrichment. PyIOmica includes optimized new algorithms adapted from MathIOmica (Mias *et al*., 2016) (which runs on the proprietary Mathematica platform), now made available as Python open source code for all users, and additionally expands extensively graphical utilities for visualization of classified temporal data, and network representation of time series. To our knowledge, there are no tools with the functionality of PyIOmica currently available in Python.

## 2 Materials and methods

### 2.1 Overview and Codebase

PyIOmica provides a complete workflow for time series processing, illustrated in the Supplementary Figure S1. The modular nature of PyIOmica allows for smooth integration with any future and existing Python tools. In addition, major data structures can be imported from MathIOmica to PyIOmica using PyIOmica functions. With PyIOmica, any results can be visualized, exported and analyzed for gene enrichment by means of a user-friendly Python interface.

PyIOmica’s codebase is a single Python module containing multiple groups of functions designed for annotations and enumerations, pre- and post-processing, clustering-related purposes, visualizations (heatmaps and classifications), normal and horizontal visibility graphs generation, and other core and utility components. Installation is simply performed using `pip install pyiomica`, and package dependencies are automatically addressed directly from PyPI (Python package index). Function documentation is embedded in the module, and is easily accessible at runtime (see Supplementary Material).

### 2.2 Data structure and analyses

We utilize the fast, robust, and versatile functionality of Python Pandas’ DataFrame, a two-dimensional size-mutable data structure that allows for multi-level hierarchical indexing (MultiIndex) along both axes (rows and columns). Pyiomica uses this multi-level hierarchical indexing to store metadata. For example, gene-specific information in a transcriptome dataset can be encoded by adding new row MultiIndex levels, and sample-specific information can be added using column MultiIndex levels. For storage and exchange, data are written using the Hierarchical Data Format (HDF5), which is essential when large amounts of data are supplied. The HDF5 files are easily accessed using Pandas HDFStore, which utilizes PyTables, or using h5py to read/write NumPy arrays.

An extensive set of PyIOmica pre-processing functions enables filtering low-quality signals, tagging missing or low values, normalization, standardization, merging and comparison of the datasets. The post-processing functions, such as temporal trends classification of power spectrum and spikes, are built on using the SciPy and scikit-learn Python toolkits. Additional functionality includes Gene Ontology (GO) and KEGG (Kyoto Encyclopedia of Genes and Genomes) pathway enrichment analyses for both non-temporal data, as well as for clusters identified through the automated time-series classification.

Temporal trends are automatically discovered using a power spectrum calculation based on a Lomb-Scargle transformation based algorithm(Mias *et al*., 2016), which properly accounts for missing points and/or unevenly-sampled data. Autocorrelations and periodograms are calculated and signals showing statistically significant trends are retained for downstream analysis.

### 2.3 Visibility graphs and classification visualization

PyIOmica introduces a Python implementation of the standard normal and horizontal visibility graphs(Lacasa *et al*., 2008), whose adjacency matrices are calculated using two methods: a Numba JIT-accelerated approach scalable to multiple CPUs, and a NumPy-accelerated approach, which is preferred for large datasets but not readily scalable beyond a few CPUs. The adjacency matrix is then converted to a NetworkX graphs for visualization and analysis.

We used Python Matplotlib plotting functions to visualize histograms, dendrograms, heatmaps and visibility graphs on different layouts. Figure 1(a) shows example RNA-sequencing gene expression data from a 24 hour time-series, clustered into two groups based on autocorrelations of the gene expression. Subgroups are then determined from the gene expression in each autocorrelation group. The data from group 1, subgroup 2 containing 191 genes is visualised in Figure 1(b) and (c) as a visibility graph on a circular and linear layout, respectively. Temporal events are detected and indicated with solid blue lines encompassing groups of points, or communities. Additional examples are provided as a Jupyter notebook (Supplementary Material, using data that is provided with the PyIOmica software release).

**Fig. 1.**
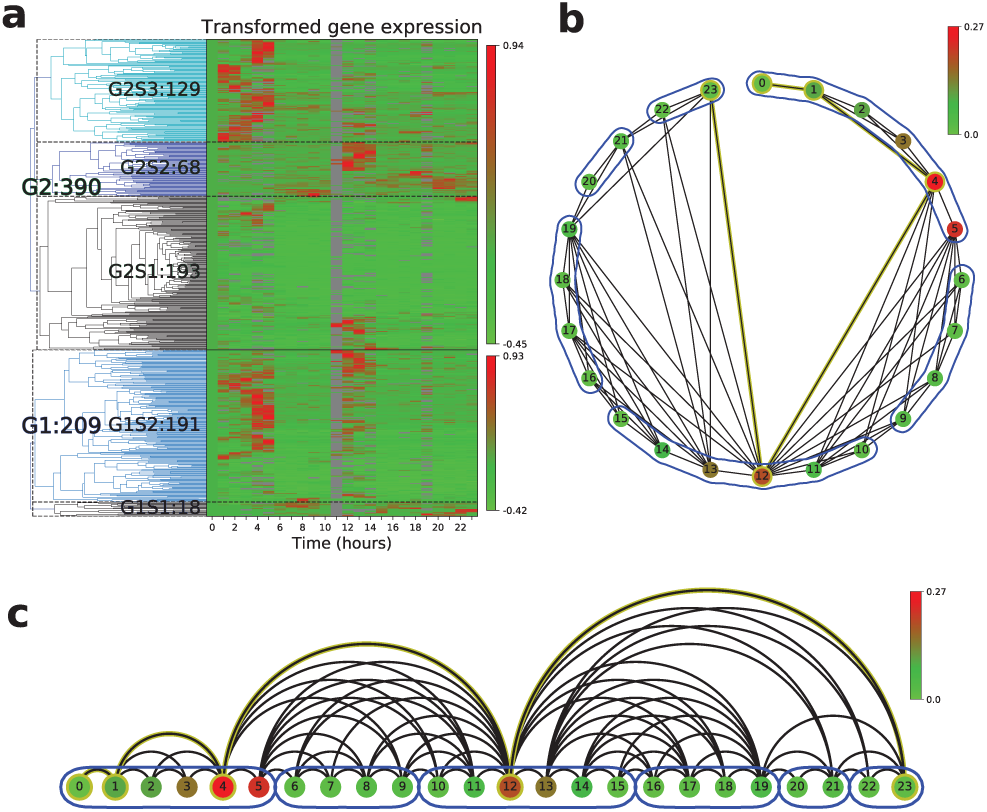
Example of PyIOImica data visualization. a Dendrogram with heatmap of automatically classified longitudinal gene expression data. Autocorrelations are used to identify temporal trends in the data. Sub-groups are determined based on similar collective behavior over time. b Visibility graph of the median signal intensity extracted from group G1S2, panel a, on a circular layout. c Same graph visualization as in b on linear layout.

## 3 Conclusion

The open source PyIOmica Python package characterizes time-series from multiple omics and classify temporal trends with a streamlined automated pipeline based on spectral analysis. PyIOmica also offers broad bioinformatics functionality, including clustering, visualization, and enrichment, and extends previous developments (Mias *et al*., 2016) to an open-source, community-accessible platform for data science. We anticipate future versions of PyIOmica to utilize its codebase flexibility to expand its bioinformatics tools for genomic as well as differential gene expresion analyses, and graph construction and characterization.

