## Supplementary Material for "PyIOmica: Longitudinal Omics Analysis and Classification"

#### 1 SUPPLEMENTARY TEXT

##### 1.1 PyIOmica information

PyIOmica is a Python package released under an MIT license through the Python Package Index. The current version used in this manuscript is 1.0.0 (07/19/19).

##### 1.2 PyIOmica Installation Instructions

The following installation instructions have been tested on Mac OS 10.14.5, Windows 10, and Ubuntu using the Anaconda Python Distribution.

###### 1.2.1 Pre-Installation Requirements

To install PyIOmica on any platform you need:

- Python version 3.7 or higher
- Recommended: 16GB of RAM or higher

###### 1.2.2 Installation

To install the current release from PyPI (Python Package Index) use pip:

```
$ pip install pyiomica
```

Alternatively, you can install directly from github using:

```
$ pip install git+https://github.com/gmiaslab/pyiomica/
```

##### 1.3 Running PyIOmica

After installation you can run PyIOmica in the Python REPL:

```
>>> from pyiomica import pyiomica
```

##### 1.4 Documentation

Documentation for PyIOmica is built-in and is available through the help command, for the complete documentation:

```
>>> help(pyiomica)
```

Or per function. For example:

```
>>> help(function)
```

The per-function documentation is also available in the PyIOmica Manual (see Supplementary Files). Additional examples are available as a Jupyter notebook. The documentation is provided in the `docs` directory distributed with the PyIOmica release - see GitHub and Zenodo for latest versions.

#### 2 SUPPLEMENTARY FIGURES

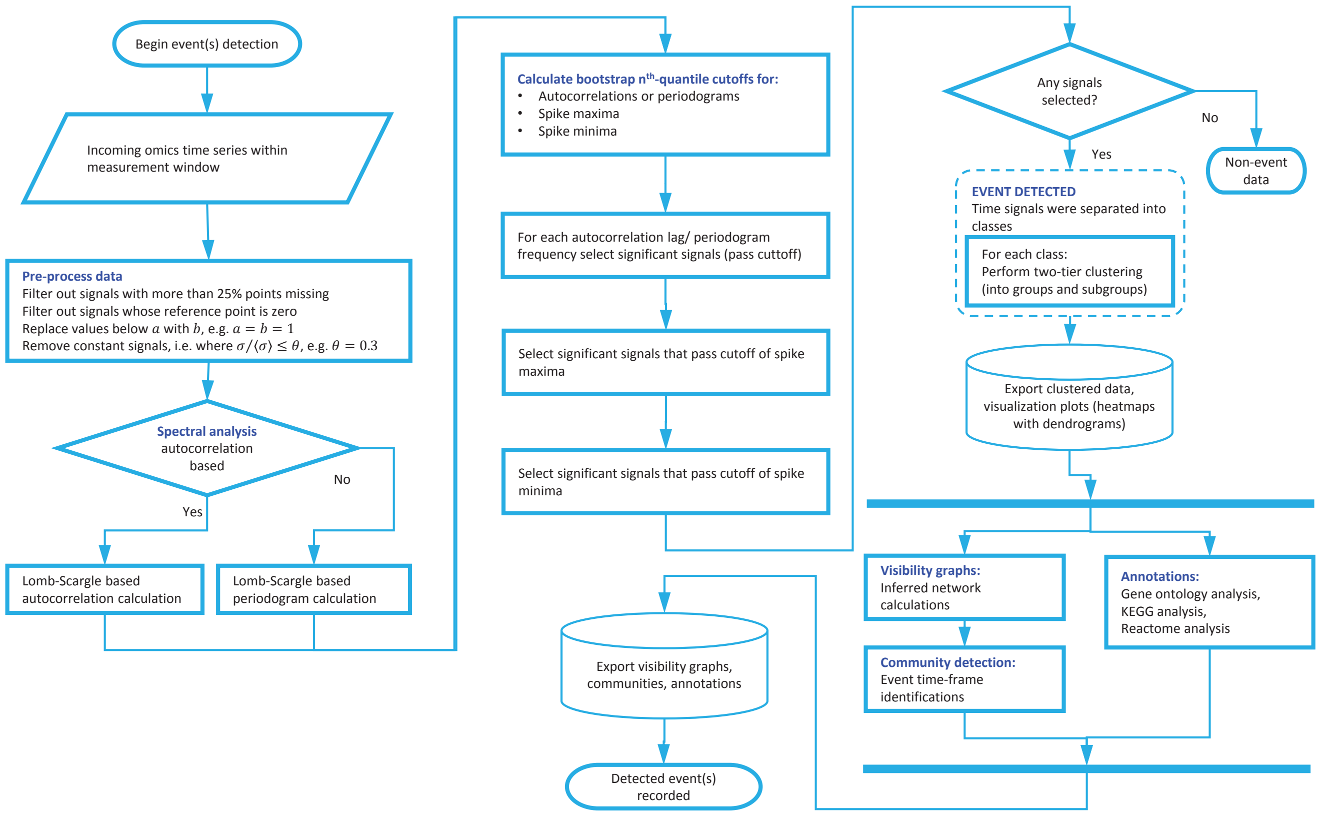


**Figure S1.** PyIOmica Scheme

### 3 PYIOMICA FUNCTION REFERENCES

#### pyiomica.pyiomica

index
PyIOmica Github Repository

```
PyIOmica is a general omics package with multiple tools for analyzing omics data.

Usage:
    from pyiomica import pyiomica
Notes:
    For additional information visit: https://github.com/gmiaslab/pyiomica and https://mathiomica.org by G. Mias Lab
```

##### Modules

| | | | |
|---|---|---|---|
| appdirs | json | matplotlib.patheffects | shutil |
| matplotlib.cm | matplotlib | pandas | sklearn |
| copy | multiprocessing | pickle | urllib |
| datetime | numpy | matplotlib.pyplot | zipfile |
| gzip | numba | pymysql | |
| h5py | networkx | importlib.resources | |
| scipy.cluster.hierarchy | os | scipy | |

##### Functions

**BenjaminiHochbergFDR**(pValues, SignificanceLevel=0.05)

```
HypothesisTesting BenjaminiHochbergFDR correction

Args:
    pValues: p-values (1D array of floats)
    SignificanceLevel: default = 0.05.

Returns:
    Corrected p-Values, p- and q-Value cuttoffs

Usage:
    result = BenjaminiHochbergFDR(pValues)
```

**ExportEnrichmentReport**(data, AppendString='', OutputDirectory=None)

```
Export results from enrichment analysis to Excel spreadsheets.

Args:
    data: enrichment results
    AppendString: custom report name, if empty then time stamp will be used
    OutputDirectory: path of directories where the report will be saved

Returns:
    None

Usage:
    ExportEnrichmentReport(goExample1, AppendString='goExample1', OutputDirectory=None)
```

**GOAnalysis**(data, GetGeneDictionaryOptions={}, AugmentDictionary=True, InputID=['UniProt ID', 'Gene Symbol'], OutputID='UniProt ID', GOAnalysisAssignerOpti[illegible] Species='human', OntologyLengthFilter=2, ReportFilter=1, ReportFilterFunction=<ufunc 'greater_equal'>, pValueCutoff=0.05, TestFunction=<function <lambda> at 0x7[illegible] HypothesisFunction=<function <lambda> at 0x7fad40c80c80>, FilterSignificant=True, OBODictionaryVariable=None, OBOGODictionaryOptions={}, MultipleListCor[illegible] MultipleList=False, GeneDictionary=None)

```
Calculate input data over-representation analysis for Gene Ontology (GO) categories.

Args:
    data: data to analyze
    GetGeneDictionaryOptions: a list of options that will be passed to this internal GetGeneDictionary function
    AugmentDictionary: a choice whether or not to augment the current ConstantGeneDictionary global variable or create a new one
    InputID:  kind of identifiers/accessions used as input
    OutputID: kind of IDs/accessions to convert the input IDs/accession numbers in the function's analysis
    GOAnalysisAssignerOptions: a list of options that will be passed to the internal GOAnalysisAssigner function
    BackgroundSet: background list to create annotation projection to limited background space, involves
considering pathways/groups/sets and that provides a list of IDs (e.g. gene accessions) that should be
considered as the background for the calculation
    Species: the species considered in the calculation, by default corresponding to human
    OntologyLengthFilter: function that can be used to set the value for which terms to consider in the computation,
by excluding GO terms that have fewer items compared to the OntologyLengthFilter value. It is used by the internal
GOAnalysisAssigner function
    ReportFilter: functions that use pathways/ontologies/groups, and provides a cutoff for membership in ontologies/pathways/groups
in selecting which terms/categories to return. It is typically used in conjunction with ReportFilterFunction
    ReportFilterFunction: specifies what operator form will be used to compare against ReportFilter option value in
selecting which terms/categories to return
    HypothesisFunction: allows the choice of function for implementing multiple hypothesis testing considerations
    FilterSignificant: can be set to True to filter data based on whether the analysis result is statistically significant,
or if set to False to return all membership computations
    OBODictionaryVariable: a GO annotation variable. If set to None, OBOGODictionary will be used internally to
automatically generate the default GO annotation
    OBOGODictionaryOptions: a list of options to be passed to the internal OBOGODictionary function that provides the GO annotations
    MultipleListCorrection: specifies whether or not to correct for multi-omics analysis. The choices are None, Automatic,
or a custom number, e.g protein+RNA
    MultipleList: specifies whether the input accessions list constituted a multi-omics list input that is annotated so
    GeneDictionary: points to an existing variable to use as a gene dictionary in annotations. If set to None
the default ConstantGeneDictionary will be used

Returns:
    Enrichment dictionary

Usage:
```

### 4 PYIOMICA FUNCTION REFERENCES

```
        goExample1 = GOAnalysis(["TAB1", "TNFSF13B", "MALT1", "TIRAP", "CHUK",
                                "TNFRSF13C", "PARP1", "CSNK2A1", "CSNK2A2", "CSNK2B", "LTBR",
                                "LYN", "MYD88", "GADD45B", "ATM", "NFKB1", "NFKB2", "NFKBIA",
                                "IRAK4", "PIAS4", "PLAU"])
```

**GOAnalysisAssigner**(PyIOmicaDataDirectory=None, ImportDirectly=False, BackgroundSet=[], Species='human', LengthFilter=None, LengthFilterFunction=<ufunc 'g
GOFileName=None, GOFileColumns=[2, 5], GOURL='http://current.geneontology.org/annotations/')

```
    Download and create gene associations and restrict to required background set.

    Args:
        PyIOmicaDataDirectory: the directory where the default package data is stored
        ImportDirectly: import from URL regardles is the file already exists
        BackgroundSet: background list to create annotation projection to limited background space, involves
    considering pathways/groups/sets and that provides a list of IDs (e.g. gene accessions) that should
    be considered as the background for the calculation
        Species: species considered in the calculation, by default corresponding to human
        LengthFilterFunction: performs computations of membership in pathways/ontologies/groups/sets,
    that specifies which function to use to filter the number of members a reported category has
    compared to the number typically provided by LengthFilter
        LengthFilter: argument for LengthFilterFunction
        GOFileName: the name for the specific GO file to download from the GOURL if option ImportDirectly is set to True
        GOFileColumns: columns to use for IDs and GO:accessions respectively from the downloaded GO annotation file,
    used when ImportDirectly is set to True to obtain a new GO association file
        GOURL: the location (base URL) where the GO association annotation files are downloaded from

    Returns:
        IDToGO and GOToID dictionary

    Usage:
        GOassignment = GOAnalysisAssigner()
```

**GeneTranslation**(InputList, TargetIDList, GeneDictionary, InputID=None, Species='human')

```
    Use geneDictionary to convert inputList IDs to different annotations as indicated by targetIDList.

    Args:
        InputList: list of names
        TargetIDList: target ID list
        GeneDictionary: an existing variable to use as a gene dictionary in annotations.
    If set to None the default ConstantGeneDictionary will be used
        InputID: the kind of identifiers/accessions used as input
        Species: the species considered in the calculation, by default corresponding to human

    Returns:
        Dictionary

    Usage:
        GenDict = GeneTranslation(data, "UniProt ID", ConstantGeneDictionary, InputID = ["UniProt ID","Gene Symbol"],  Species = "human")
```

**GetGeneDictionary**(geneUCSCTable=None, UCSCSQLString=None, UCSCSQLSelectLabels=None, ImportDirectly=False, Species='human', KEGGUCSCSplit=[Tru

```
    Create an ID/accession dictionary from a UCSC search - typically of gene annotations.

    Args:
        geneUCSCTable: path to a geneUCSCTable file
        UCSCSQLString: an association to be used to obtain data from the UCSC Browser tables. The key of the association must
    match the Species option value used (default: human). The value for the species corresponds to the actual MySQL command used
        UCSCSQLSelectLabels: an association to be used to assign key labels for the data improted from the UCSC Browser tables.
    The key of the association must match the Species option value used (default: human). The value is a multi component string
    list corresponding to the matrices in the data file, or the tables used in the MySQL query provided by UCSCSQLString
        ImportDirectly: import from URL regardles is the file already exists
        Species: species considered in the calculation, by default corresponding to human
        KEGGUCSCSplit: a two component list, {True/False, label}. If the first component is set to True the initially imported KEGG IDs,
    identified by the second component label,  are split on + string to fix nomenclature issues, retaining the string following +

    Returns:
        Dictionary

    Usage:
        geneDict = GetGeneDictionary()
```

**KEGGAnalysis**(data, AnalysisType='Genomic', GetGeneDictionaryOptions={}, AugmentDictionary=True, InputID=['UniProt ID', 'Gene Symbol'], OutputID='KEGG G
['cpd'], MolecularOutputID='cpd', KEGGAnalysisAssignerOptions={}, BackgroundSet=[], KEGGOrganism='hsa', KEGGMolecular='cpd', KEGGDatabase='pathway', P
ReportFilter=1, ReportFilterFunction=<ufunc 'greater_equal'>, pValueCutoff=0.05, TestFunction=<function <lambda> at 0x7fad40c80f28>, HypothesisFunction=<funct
0x7fad40c81048>, FilterSignificant=True, KEGGDictionaryVariable=None, KEGGDictionaryOptions={}, MultipleListCorrection=None, MultipleList=False, GeneDicti
Species='human', MolecularSpecies='compound', NonUCSC=False, PyIOmicaDataDirectory=None)

```
    Calculate input data over-representation analysis for KEGG: Kyoto Encyclopedia of Genes and Genomes pathways.
    Input can be a list, a dictionary of lists or a clustering object.

    Args:
        data: data to analyze
        AnalysisType: analysis methods that may be used, "Genomic", "Molecular" or "All"
        GetGeneDictionaryOptions: a list of options that will be passed to this internal GetGeneDictionary function
        AugmentDictionary: a choice whether or not to augment the current ConstantGeneDictionary global variable or create a new one
        InputID: the kind of identifiers/accessions used as input
        OutputID: a string value that specifies what kind of IDs/accessions to convert the input IDs/accession
    numbers in the function's analysis
        MolecularInputID: a string list to indicate the kind of ID to use for the input molecule entries
        KEGGAnalysisAssignerOptions: a list of options that will be passed to this internal KEGGAnalysisAssigner function
        BackgroundSet: a list of IDs (e.g. gene accessions) that should be considered as the background for the calculation
        KEGGOrganism: indicates which organism (org) to use for "Genomic" type of analysis (default is human analysis: org="hsa")
        KEGGMolecular: which database to use for molecular analysis (default is the compound database: cpd)
        KEGGDatabase: KEGG database to use as the target database
        PathwayLengthFilter: pathways to consider in the computation, by excluding pathways that have fewer items
    compared to the PathwayLengthFilter value
        ReportFilter: provides a cutoff for membership in ontologies/pathways/groups in selecting which terms/categories
    to return. It is typically used in conjunction with ReportFilterFunction
```

### 5 PYIOMICA FUNCTION REFERENCES

```
        ReportFilterFunction: operator form will be used to compare against ReportFilter option value in selecting
    which terms/categories to return
        pValueCutoff: a cutoff p-value for (adjusted) p-values to assess statistical significance
        TestFunction: a function used to calculate p-values
        HypothesisFunction: allows the choice of function for implementing multiple hypothesis testing considerations
        FilterSignificant: can be set to True to filter data based on whether the analysis result is statistically significant,
    or if set to False to return all membership computations
        KEGGDictionaryVariable: KEGG dictionary, and provides a KEGG annotation variable. If set to None, KEGGDictionary
    will be used internally to automatically generate the default KEGG annotation
        KEGGDictionaryOptions: a list of options to be passed to the internal KEGGDictionary function that provides the KEGG annotations
        MultipleListCorrection: specifies whether or not to correct for multi-omics analysis.
    The choices are None, Automatic, or a custom number
        MultipleList: whether the input accessions list constituted a multi-omics list input that is annotated so
        GeneDictionary: existing variable to use as a gene dictionary in annotations. If set to None the default ConstantGeneDictionary wi
        Species: the species considered in the calculation, by default corresponding to human
        MolecularSpecies: the kind of molecular input
        NonUCSC: if UCSC browser was used in determining an internal GeneDictionary used in ID translations,
    where the KEGG identifiers for genes are number strings (e.g. 4790).The NonUCSC option can be set to True
    if standard KEGG accessions are used in a user provided GeneDictionary variable,
    in the form OptionValue[KEGGOrganism] <>:<>numberString, e.g. hsa:4790
        PyIOmicaDataDirectory: directory where the default package data is stored

    Returns:
        Enrichment dictionary

    Usage:
        keggExample1 = KEGGAnalysis(["TAB1", "TNFSF13B", "MALT1", "TIRAP", "CHUK", "TNFRSF13C", "PARP1", "CSNK2A1", "CSNK2A2", "CSNK2B", "
                                    "GADD45B", "ATM", "NFKB1", "NFKB2", "NFKBIA", "IRAK4", "PIAS4", "PLAU", "POLR3B", "NME1", "CTP
```

**KEGGAnalysisAssigner**(PyIOmicaDataDirectory=None, ImportDirectly=False, BackgroundSet=[], KEGGQuery1='pathway', KEGGQuery2='hsa', LengthFilter=None,
<ufunc 'greater_equal'>, Labels=['IDToPath', 'PathToID'])

```
    Create KEGG: Kyoto Encyclopedia of Genes and Genomes pathway associations,
    restricted to required background set, downloading the data if necessary.

    Args:
        PyIOmicaDataDirectory: directory where the default package data is stored
        ImportDirectly: import from URL regardles is the file already exists
        BackgroundSet: a list of IDs (e.g. gene accessions) that should be considered as the background for the calculation
        KEGGQuery1: make KEGG API calls, and sets string query1 in http://rest.kegg.jp/link/<> query1 <> / <> query2.
    Typically this will be used as the target database to find related entries by using database cross-references
        KEGGQuery2: KEGG API calls, and sets string query2 in http://rest.kegg.jp/link/<> query1 <> / <> query2.
    Typically this will be used as the source database to find related entries by using database cross-references
        LengthFilterFunction: option for functions that perform computations of membership in
    pathways/ontologies/groups/sets, that specifies which function to use to filter the number of members a reported
    category has compared to the number typically provided by LengthFilter
        LengthFilter: allows the selection of how many members each category can have, as typically
    restricted by the LengthFilterFunction
        Labels: a string list for how keys in a created association will be named

    Returns:
        IDToPath and PathToID dictionary

    Usage:
        KEGGassignment = KEGGAnalysisAssigner()
```

**KEGGDictionary**(PyIOmicaDataDirectory=None, ImportDirectly=False, KEGGQuery1='pathway', KEGGQuery2='hsa')

```
    Create a dictionary from KEGG: Kyoto Encyclopedia of Genes and Genomes terms -
    typically association of pathways and members therein.

    Args:
        PyIOmicaDataDirectory: directory where the default package data is stored
        ImportDirectly: import from URL regardles is the file already exists
        KEGGQuery1: make KEGG API calls, and sets string query1 in http://rest.kegg.jp/link/<> query1 <> / <> query2.
    Typically this will be used as the target database to find related entries by using database cross-references
        KEGGQuery2: KEGG API calls, and sets string query2 in http://rest.kegg.jp/link/<> query1 <> / <> query2.
    Typically this will be used as the source database to find related entries by using database cross-references

    Returns:
        Dictionary of definitions

    Usage:
        KEGGDict = KEGGDictionary()
```

**LombScargle**(inputTimes, inputData, inputSetTimes, FrequenciesOnly=False, NormalizeIntensities=False, OversamplingRate=1, UpperFrequencyFactor=1)

```
    Calculate Lomb-Scargle periodogram.

    Args:
        inputTimes: times corresponding to provided data points (1D array of floats)
        inputData: data points (1D array of floats)
        inputSetTimes: a complete set of all possible N times during which data could have been collected
        FrequenciesOnly: return frequencies only
        NormalizeIntensities: normalize intensities to unity
        OversamplingRate: oversampling rate
        UpperFrequencyFactor: upper frequency factor

    Returns:
        Periodogram with a list of frequencies.

    Usage:
        pgram = LombScargle(inputTimes, inputData, inputSetTimes)
```

**MassDictionary**(PyIOmicaDataDirectory=None)

```
    Load PyIOmica's current mass dictionary.

    Args:
        PyIOmicaDataDirectory: directory where the default package data is stored
```

### 6 PYIOMICA FUNCTION REFERENCES

```
Returns:
    Mass dictionary

Usage:
    MassDict = MassDictionary()
```

**MassMatcher**(data, accuracy, MassDictionaryVariable=None, MolecularSpecies='cpd')

```
Assign putative mass identification to input data based on monoisotopic mass
(using PyIOmica's mass dictionary). The accuracy in parts per million.

Args:
    data: input data
    accuracy: accuracy
    MassDictionaryVariable: mass dictionary variable. If set to None, inbuilt
mass dictionary (MassDictionary) will be loaded and used
    MolecularSpecies: the kind of molecular input

Returns:
    List of IDs

Usage:
   result = MassMatcher(18.010565, 2)
```

**OBOGODictionary**(FileURL='http://purl.obolibrary.org/obo/go/go-basic.obo', ImportDirectly=False, PyIOmicaDataDirectory=None, OBOFile='goBasicObo.txt')

```
Generate Open Biomedical Ontologies (OBO) Gene Ontology (GO) vocabulary dictionary.

Args:
    FileURL: provides the location of the Open Biomedical Ontologies (OBO) Gene Ontology (GO)
file in case this will be downloaded from the web
    ImportDirectly: import from URL regardles is the file already exists
    PyIOmicaDataDirectory: path of directories to data storage
    OBOFile: name of file to store data in (file will be zipped)

Returns:
    Dictionary of definitions

Usage:
    OBODict = OBOGODictionary()
```

**PlotHorizontalVisibilityGraph**(A, data, times, fileName, id)

```
Bar-plot style horizontal visibility graph.

Args:
    A: Adjacency matrix
    data: Numpy 2-D array of floats
    times: Numpy 1-D array of floats
    fileName: name of the figure file to save
    id: label to add to the figure title

Returns:
    None

Usage:
PlotHorizontalVisibilityGraph(A, data, times, 'FIgure.png', 'Test Data')
```

**PlotVisibilityGraph**(A, data, times, fileName, id)

```
Bar-plot style visibility graph.

Args:
    A: Adjacency matrix
    data: Numpy 2-D array of floats
    times: Numpy 1-D array of floats
    fileName: name of the figure file to save
    id: label to add to the figure title

Returns:
    None

Usage:
    PlotVisibilityGraph(A, data, times, 'FIgure.png', 'Test Data')
```

**addVisibilityGraph**(data, times, dataName='G1S1', coords=[0.05, 0.95, 0.05, 0.95], numberOfVGs=1, groups_ac_colors=['b'], fig=None, numberOfCommunities=6, pri
fontsize=None, nodesize=None, level=0.55, commLineWidth=0.5, lineWidth=1.0, withLabel=True, withTitle=False, layout='circle', radius=0.07, noplot=False)

```
Draw a Visibility graph of data on a provided Matplotlib figure.

Args:
    data: array of data to visualize
    times: times corresponding to each data point, used for labels
    dataName: label to include in file name
    coords: coordinates of location of the plot on the figure
    numberOfVGs: number of plots to add to this figure
    groups_ac_colors: colors corresponding to different groups of graphs
    fig: figure object
    printCommunities: print communities details to screen
    fontsize: size of labels
    nodesize: size of nodes
    level: distance of the community lines to nodes
    commLineWidth: width of the community lines
    lineWidth: width of the edges between nodes
    withLabel: include label on plot
    withTitle: include title on plot

Returns:
    None
```

### 7 PYIOMICA FUNCTION REFERENCES

```
    Usage:
        addVisibilityGraph(exampleData, exampleTimes, fig=fig, fontsize=16, nodesize=700,
                           level=0.85, commLineWidth=3.0, lineWidth=2.0, withLabel=False)
```

**ampSquaredNormed**(func, freq, times, data)

```
    Lomb-Scargle core function
    Calculate the different frequency components of our spectrum: project the cosine/sine component and normalize it:

    Args:
        func: Sin or Cos
        freq: frequencies (1D array of floats)
        times: input times (starting point adjusted w.r.t.dataset times), Zero-padded
        data: input Data with the mean subtracted from it, before zero-padding.

    Returns:
        Squared amplitude normalized.

    Usage:
        coef = ampSquaredNormed(np.cos, frequency, inputTimesNormed, inputDataCentered)
        Intended for internal use only.
```

**autocorrelation**(inputTimes, inputData, inputSetTimes, UpperFrequencyFactor=1)

```
    Autocorrelation function

    Args:
        inputTimes: times corresponding to provided data points (1D array of floats)
        inputData: data points (1D array of floats)
        inputSetTimes: a complete set of all possible N times during which data could have been collected.

    Returns:
        Array of time lags with corresponding autocorrelations

    Usage:
        result = autocorrelation(inputTimes, inputData, inputSetTimes)
```

**boxCoxTransform**(subset, lmbda=None, giveLmbda=False)

```
    Power transform from scipy.stats

    Args:
        subset: pandas Series.
        lmbda: Lambda parameter, if not specified optimal value will be determined
        giveLmbda: also return Lambda value

    Returns:
        Transformed subset and Lambda parameter

    Usage:
        myData = boxCoxTransform(myData)
```

**boxCoxTransformDataframe**(df)

```
    Box-cox transform data.

    Args:
        df: pandas DataFrame

    Returns:
        Processed pandas Dataframe

    Usage:
        df_data = boxCoxTransformDataframe(df_data)
```

**chop**(expr, tolerance=1e-10)

```
    Equivalent of Mathematica.Chop Function.

    Args:
        expr: a number or a pyhton sequence of numbers
        tolerance: default is the same as in Mathematica

    Returns:
        Chopped data

    Usage
        data = chop(data)
```

**compareTimeSeriesToPointDataframe**(df, point='first')

```
    Subtract a particular point of each time series (row) of a Dataframe.

    Args:
        df: pandas DataFrame
        point: 'first', 'last', 0, 1, ... , 10, or a value.

    Returns:
        Processed pandas Dataframe

    Usage:
        df_data = compareTimeSeriesToPointDataframe(df_data)
```

**compareTwoTimeSeriesDataframe**(df1, df2, function=<ufunc 'subtract'>, compareAllLevelsInIndex=True, mergeFunction=<function mean at 0x7fad301bd840>)

```
    Create a new Dataframe based on comparison of two existing Dataframes.

    Args:
        df1: pandas DataFrame
        df2: pandas DataFrame
        function: np.subtract (default), np.add, np.divide, or another <ufunc>.
        compareAllLevelsInIndex: True (default), if False only "source" and "id" will be compared,
        mergeFunction: input Dataframes are merged with this function, i.e. np.mean (default), np.median, np.max, or another <ufunc>.
```

### 8 PYIOMICA FUNCTION REFERENCES

```
    Returns:
        New merged pandas Dataframe

    Usage:
        df_data = compareTwoTimeSeriesDataframe(df_dataH2, df_dataH1, function=np.subtract, compareAllLevelsInIndex=False, mergeFunction=n
```

**createDirectories**(path)

```
    Create a path of directories, unless the path already exists.

    Args:
        path: path directory

    Returns:
        None

    Usage:
        createDirectories("/pathToFolder1/pathToSubFolder2")
```

**createReverseDictionary**(inputDictionary)

```
    Efficient way to create a reverse dictionary from a dictionary.
    Utilizes Pandas.Dataframe.groupby and Numpy arrays indexing.

    Args:
        inputDictionary: a dictionary to reverse

    Returns:
        Reversed dictionary

    Usage:
        revDict = createReverseDictionary(Dict)
```

**exportClusteringObject**(ClusteringObject, saveDir, dataName, includeData=True, includeAutocorr=True)

```
    Export a clustering Groups-Subgroups dictionary object to a SpreadSheet.
    Linkage data is not exported.

    Args:
        ClusteringObject: clustering object
        saveDir: path of directories to save the object to
        dataName: label to include in the file name
        includeData: export data
        includeAutocorr: export autocorrelations of data

    Returns:
        File name of the exported clustering object

    Usage:
        exportClusteringObject(myObj, '/dir1', 'myObj')
```

**filterOutAllZeroSignalsDataframe**(df)

```
    Filter out all-zero signals from a DataFrame.

    Args:
        df: pandas DataFrame

    Returns:
        Processed pandas Dataframe

    Usage:
        df_data = filterOutAllZeroSignalsDataframe(df_data)
```

**filterOutFirstPointZeroSignalsDataframe**(df)

```
    Filter out out first time point zeros signals from a DataFrame.

    Args:
        df: pandas DataFrame

    Returns:
        Processed pandas Dataframe

    Usage:
        df_data = filterOutFirstPointZeroSignalsDataframe(df_data)
```

**filterOutFractionZeroSignalsDataframe**(df, max_fraction_of_allowed_zeros)

```
    Filter out fraction-zero signals from a DataFrame.

    Args:
        df: pandas DataFrame
        max_fraction_of_allowed_zeros: maximum fraction of allowed zeros

    Returns:
        Processed pandas Dataframe

    Usage:
        df_data = filterOutFractionZeroSignalsDataframe(df_data, 0.75)
```

**getAdjacencyMatrixOfHorizontalVisibilityGraph**(data)

```
    Calculate adjacency matrix of horizontal visibility graph.
    JIT-accelerated version (a bit faster than NumPy-accelerated version).
    Single-threaded beats NumPy up to 2k data sizes.
    Allows use of Multiple CPUs.

    Args:
        data: Numpy 2-D array of floats

    Returns:
```

### 9 PYIOMICA FUNCTION REFERENCES

```
        Adjacency matrix

    Usage:
        A = getAdjacencyMatrixOfHorizontalVisibilityGraph(data)
```

**getAdjacencyMatrixOfHorizontalVisibilityGraph_NUMPY**(data)

```
    Calculate adjacency matrix of horizontal visibility graph.
    NumPy-accelerated version.
    Use with datasets larger than 2k.
    Use in serial applications.

    Args:
        data: Numpy 2-D array of floats

    Returns:
        Adjacency matrix

    Usage:
        A = getAdjacencyMatrixOfHorizontalVisibilityGraph_NUMPY(data)
```

**getAdjacencyMatrixOfVisibilityGraph**(data, times)

```
    Calculate adjacency matrix of visibility graph.
    JIT-accelerated version (a bit faster than NumPy-accelerated version).
    Allows use of Multiple CPUs.

    Args:
        data: Numpy 2-D array of floats
        times: Numpy 1-D array of floats

    Returns:
        Adjacency matrix

    Usage:
        A = getAdjacencyMatrixOfVisibilityGraph_serial(data, times)
```

**getAdjacencyMatrixOfVisibilityGraph_NUMPY**(data, times)

```
    Calculate adjacency matrix of visibility graph.
    NumPy-accelerated version. Somewhat slower than JIT-accelerated version.
    Use in serial applications.

    Args:
        data: Numpy 2-D array of floats
        times: Numpy 1-D array of floats

    Returns:
        Adjacency matrix

    Usage:
        A = getAdjacencyMatrixOfVisibilityGraph_serial(data, times)
```

**getAutocorrelationsOfData**(params)

```
    Calculate autocorrelation using Lomb-Scargle Autocorrelation.
    NOTE: there should be already no missing or non-numeric points in the input Series or Dataframe

    Args:
        params: a tuple of parameters in the form (df_data, setAllInputTimes), where
        df_data is a pandas Series or Dataframe,
        setAllInputTimes is a complete set of all possible N times during which data could have been collected.

    Returns:
        Array of autocorrelations of data.

    Usage:
        result  = autocorrelation(df_data, setAllInputTimes)
```

**getEstimatedNumberOfClusters**(data, cluster_num_min, cluster_num_max, trials_to_do, numberOfAvailableCPUs=4, plotID=None, printScores=False)

```
    Get estimated number of clusters using ARI with KMeans

    Args:
        data: data to analyze
        cluster_num_min: minimum possible number of clusters
        cluster_num_max: maximum possible number of clusters
        trials_to_do: number of trials to do in ARI function
        numberOfAvailableCPUs: number of processes to run in parallel
        plotID: label for the plot of peaks
        printScores: print all scores

    Returns:
        Largest peak, other possible peaks.

    Usage:
        n_clusters = getEstimatedNumberOfClusters(data, 1, 20, 25)
```

**getGroupingIndex**(data, n_groups=None, method='weighted', metric='correlation', significance='Elbow')

```
    Cluster data into N groups, if N is provided, else determine N
    return: linkage matrix, cluster labels, possible cluster labels.

    Args:
        data: data to analyze
        n_groups: number of groups to split data into
        method: linkage calculation method
        metric: distance measure
        significance: method for determining optimal number of groups and subgroups

    Returns:
        Linkage matrix, cluster index, possible groups
```

### 10 PYIOMICA FUNCTION REFERENCES

```
Usage:
    x, y, z = getGroupingIndex(data, method='weighted', metric='correlation', significance='Elbow')
```

**getLobmScarglePeriodogramOfDataframe**(df_data, NumberOfCPUs=4, parallel=True)

```
Calculate Lobm-Scargle periodogram of DataFrame.

Args:
    df: pandas DataFrame
    parallel: calculate in parallel mode (>1 process)
    NumberOfCPUs: number of processes to create if parallel

Returns:
    New pandas Dataframe

Usage:
    df_periodograms = getLobmScarglePeriodogramOfDataframe(df_data)
```

**getRandomAutocorrelations**(df_data, NumberOfRandomSamples=100000, NumberOfCPUs=4)

```
Generate autocorrelation null-distribution from permutated data using Lomb-Scargle Autocorrelation.
NOTE: there should be already no missing or non-numeric points in the input Series or Dataframe

Args:
    df_data: pandas Series or Dataframe
    NumberOfRandomSamples: size of the distribution to generate
    NumberOfCPUs: number of processes to run simultaneously

Returns:
    DataFrame containing autocorrelations of null-distribution of data.

Usage:
    result = getRandomAutocorrelations(df_data)
```

**getRandomPeriodograms**(df_data, NumberOfRandomSamples=100000, NumberOfCPUs=4)

```
Generate periodograms null-distribution from permutated data using Lomb-Scargle function.

Args:
    df_data: pandas Series or Dataframe
    NumberOfRandomSamples: size of the distribution to generate
    NumberOfCPUs: number of processes to run simultaneously

Returns:
    New Pandas DataFrame containing periodograms

Usage:
    result = getRandomPeriodograms(df_data)
```

**getSpikes**(inputData, func, cutoffs)

```
Get sorted index of signals with statistically significant spikes,
i.e. those that pass the provided cutoff.

Args:
    inputData: data points (2D array of floats) where rows are normalized signals
    func: np.max or np.min
    cutoffs: a dictionary of cutoff values

Returns:
    Index of data with statistically significant spikes

Usage:
    index = getSpikes(inputData, np.max, cutoffs)
```

**getSpikesCutoffs**(df_data, p_cutoff, NumberOfRandomSamples=1000)

```
Calculate spikes cuttoffs from a bootstrap of provided data,
gived the significance cutoff p_cutoff.

Args:
    df_data: pandas DataFrame where rows are normalized signals
    p_cutoff: p-Value cutoff, e.g. 0.01
    NumberOfRandomSamples: size of the bootstrap distribution

Returns:
    Dictionary of spike cutoffs.

Usage:
    cutoffs = getSpikesCutoffs(df_data, 0.01)
```

**get_optimal_number_clusters_from_linkage_Elbow**(Y)

```
Get optimal number clusters from linkage.
A point of the highest accelleration of the fusion coefficient of the given linkage.

Args:
    Y: linkage matrix

Returns:
    Optimal number of clusters

Usage:
    n_clusters = get_optimal_number_clusters_from_linkage_Elbow(Y)
```

**get_optimal_number_clusters_from_linkage_Silhouette**(Y, data, metric)

```
Determine the optimal number of cluster in data maximizing the Silhouette score.

Args:
    Y: linkage matrix
    data: data to analyze
```

### 11 PYIOMICA FUNCTION REFERENCES

```
    metric: distance measure

Returns:
    Optimal number of clusters

Usage:
    n_clusters = get_optimal_number_clusters_from_linkage_Elbow(Y, data, 'euclidean')
```

**hdf5_usage_information**()

```
Store/export any lagge datasets in hdf5 format via 'pandas' or 'h5py'

# mode='w' creates/recreates file from scratch
# mode='a' creates (if no file exists) or appends to the existing file, and reads it
# mode='r' is read only

# Save data to file using 'pandas':
df_example = pd.DataFrame({'A': [1, 2, 3], 'B': [4, 5, 6]}, index=['a', 'b', 'c'])
df_example.to_hdf('data.h5', key='my_df1', mode='a')
or
series_example = pd.Series([1, 2, 3, 4])
series_example.to_hdf('data.h5', key='my_series', mode='a')

# Create groups and datasets using 'h5py' and 'numpy' arrays:
tempFile = h5py.File('data.h5', 'a')
tempArray = np.array([[1,2,3,4,5],[6,7,8,9,10]]).astype(float)
if not 'arrays/my_array' in tempFile:
    dataset_example = tempFile.create_dataset('arrays/my_array', data=tempArray, maxshape=(None,2), dtype=tempArray.dtype,
                                              chunks=True) #auto-chunked, else use e.g. chunks=(100, 2)
                                              #compression='gzip', compression_opts=6
else:
    dataset_example = tempFile['arrays/my_array']

group_example = tempFile.create_group('more_data/additional')

# Modify values by slicing the dataset or replacing etire one using [...]
dataset_example[:] = np.array([[10,2,3,4,1],[60,7,8,9,1]])

# New shapes cannot be broadcasted, the dataset needs to be resized explicitly
dataset_example.resize(dataset_example.shape[0]+10, axis=0) #add more rows (initiated with zeros)


# Read data from h5 file:
df_example = pd.read_hdf('data.h5', 'my_df1')

tempFile = h5py.File('data.h5', 'r')
array_example = tempFile['arrays/my_array'].value
```

**internalAnalysisFunction**(data, multiCorr, MultipleList, OutputID, InputID, Species, totalMembers, pValueCutoff, ReportFilterFunction, ReportFilter, TestFunction, Hy[illegible] FilterSignificant, AssignmentForwardDictionary, AssignmentReverseDictionary, prefix, infoDict)

```
Analysis for Multi-Omics or Single-Omics input list
The function is used internally and not intended to be used directly by user.

Usage:
    Intended for internal use
```

**makeClusteringObject**(df_data, df_data_autocorr, significance='Elbow')

```
Make a clustering Groups-Subgroups dictionary object.

Args:
    df_data: data to analyze in DataFrame format
    df_data_autocorr: autocorrelations or periodograms in DataFrame format
    significance: method for determining optimal number of groups and subgroups

Returns:
    Clustering object

Usage:
    myObj = makeClusteringObject(df_data, df_data_autocorr, significance='Elbow')
```

**makeDataHistograms**(df, saveDir, dataName)

```
Make a histogram for each pandas Series (time point) in a pandas Dataframe.

Args:
    df: DataFrame containing data to visualize
    saveDir: path of directories to save the object to
    dataName: label to include in the file name

Returns:
    None

Usage:
    makeDataHistograms(df, '/dir1', 'myData')
```

**makeDendrogramHeatmap**(ClusteringObject, saveDir, dataName, AutocorrNotPeriodogr=True, textScale=1.0, vectorImage=True)

```
Make Dendrogram-Heatmap plot along with VIsibility graphs.

Args:
    ClusteringObject: clustering object
    saveDir: path of directories to save the object to
    dataName: label to include in the file name
    AutocorrNotPeriodogr: export data
    textScale: scaling of text size
    vectorImage: Boolean for exporting vector graphics or PNG format

Returns:
    None
```

### 12 PYIOMICA FUNCTION REFERENCES

```
    Usage:
        makeDendrogramHeatmap(myObj, '/dir1', 'myData', AutocorrNotPeriodogr=True)
```

**makeLombScarglePeriodograms**(df, saveDir, dataName)

```
    Make a combined plot of the signal and its Lomb-Scargle periodogram
    for each pandas Series (time point) in a pandas Dataframe.

    Args:
        df: DataFrame containing data to visualize
        saveDir: path of directories to save the object to
        dataName: label to include in the file name

    Returns:
        None

    Usage:
        makeLombScarglePeriodograms(df, '/dir1', 'myData')
```

**mergeDataframes**(listOfDataframes)

```
    Merge a list of Dataframes (outer join).

    Args:
        listOfDataframes: list of pandas DataFrames

    Returns:
        New pandas Dataframe

    Usage:
        df_data = mergeDataframes([df_data1, df_data2])
```

**metricCommonEuclidean**(u, v)

```
    Metric to calculate 'euclidean' distance between vectors u and v
    using only common non-missing points (not NaNs).

    Args:
        u: Numpy 1-D array
        v: Numpy 1-D array

    Returns:
        Measure of the distance between u and v

    Usage:
        dist = metricCommonEuclidean(u,v)
```

**modifiedZScore**(subset)

```
    Calculate modified z-score of a 1D array based on "Median absolute deviation".
    Use on 1-D arrays only.

    Args:
        subset: data to transform

    Returns:
        Transformed subset

    Usage:
        data = modifiedZScore(data)
```

**modifiedZScoreDataframe**(df)

```
    Z-score (Median-based) transform data.

    Args:
        df: pandas DataFrame

    Returns:
        Processed pandas Dataframe

    Usage:
        df_data = modifiedZScoreDataframe(df_data)
```

**normalizeSignalsToUnityDataframe**(df)

```
    Normalize signals to unity.

    Args:
        df: pandas DataFrame

    Returns:
        Processed pandas Dataframe

    Usage:
        df_data = normalizeSignalsToUnityDataframe(df_data)
```

**obtainConstantGeneDictionary**(GeneDictionary, GetGeneDictionaryOptions, AugmentDictionary)

```
    Obtain gene dictionary - if it exists can either augment with new information or Species or create new,
    if not exist then create variable.

    Args:
        GeneDictionary: an existing variable to use as a gene dictionary in annotations.
    If set to None the default ConstantGeneDictionary will be used
        GetGeneDictionaryOptions: a list of options that will be passed to this internal GetGeneDictionary function
        AugmentDictionary: a choice whether or not to augment the current ConstantGeneDictionary global variable or create a new one

    Returns:
        None

    Usage:
```

### 13 PYIOMICA FUNCTION REFERENCES

```
        obtainConstantGeneDictionary(None, {}, False)
```

**pAutocorrelation**(args)

```
Wrapper of Autocorrelation function for use with Multiprocessing.

Args:
    args: a tuple of arguments in the form (inputTimes, inputData, inputSetTimes)

Returns:
    Array of time lags with corresponding autocorrelations

Usage:
    result = pAutocorrelation((inputTimes, inputData, inputSetTimes))
```

**pLombScargle**(args)

```
Wrapper of LombScargle function for use with Multiprocessing.

Args:
    args: a tuple of arguments in the form (inputTimes, inputData, inputSetTimes)

Returns:
    Array of frequencies with corresponding intensities

Usage:
    result = pLombScargle((inputTimes, inputData, inputSetTimes))
```

**prepareDataframe**(dataDir, dataFileName, AlltimesFileName)

```
Make a DataFrame from CSV files.

Args:
    dataDir: path of directories pointing to data
    dataFileName: file name in dataDir
    AlltimesFileName: file name in dataDir

Returns:
    Pandas Dataframe

Usage:
    df_data = prepareDataframe(dataDir, dataFileName, AlltimesFileName)
    df_data.index = pd.MultiIndex.from_tuples([(item.split(':')[1], item.split(':')[0].split('_')[0],
                                               (' '.join(item.split(':')[0].split('_')[1:]),)) for item in df_data.index.values],
                                               names=['source', 'id', 'metadata'])
```

**quantileNormalizeDataframe**(df)

```
Quantile Normalize signals to normal distribution.

Args:
    df: pandas DataFrame

Returns:
    Processed pandas Dataframe

Usage:
    df_data = quantileNormalizeDataframe(df_data)
```

**read**(fileName, withPKLZextension=True, hdf5fileName=None, jsonFormat=False)

```
Read object from a file recorded by function "write". Pandas and Numpy objects are
read from HDF5 file when provided, otherwise attempt to read from PKLZ file.

Args:
    fileName: path of directories ending with the file name
    withPKLZextension: add ".pklz" to a pickle file
    hdf5fileName: path of directories ending with the file name. If None then data is pickled
    jsonFormat: save data into compressed json file

Returns:
    data: data object to write into a file

Usage:
    exampleDataFrame = read('/dir1/exampleDataFrame', hdf5fileName='/dir2/data.h5')
```

**readMathIOmicaData**(fileName)

```
Read text files exported by MathIOmica and convert to Python data

Args:
    fileName: path of directories and name of the file containing data

Returns:
    Python data

Usage:
    data = readMathIOmicaData("../../MathIOmica/MathIOmica/MathIOmicaData/ExampleData/rnaExample")
```

**removeConstantSignalsDataframe**(df, theta_cutoff)

```
Remove constant signals.

Args:
    df: pandas DataFrame
    theta_cutoff: parameter for filtering the signals

Returns:
    Processed pandas Dataframe

Usage:
    df_data = removeConstantSignalsDataframe(df_data, 0.3)
```

### 14 PYIOMICA FUNCTION REFERENCES

**runCPUs**(NumberOfAvailableCPUs, func, list_of_tuples_of_func_params)

```
Parallelize function call with multiprocessing.Pool.

Args:
    NumberOfAvailableCPUs: number of processes to create
    func: function to apply, must take at most one argument
    list_of_tuples_of_func_params: function parameters

Returns:
    Results of func in a numpy array

Usage:
    results = runCPUs(4, pAutocorrelation, [(times[i], data[i], allTimes) for i in range(10)])
```

**runForClusterNum**(arguments)

```
Calculate Adjusted Rand Index of the data for a range of cluster numbers.

Args:
    arguments: a tuple of three parameters int the form
    (cluster_num, data_array, trials_to_do), where
    cluster_num: maximum number of clusters
    data_array: data to test
    trials_to_do: number of trials for each cluster number

Returns:
    Numpy array

Usage:
    instPool = multiprocessing.Pool(processes = NumberOfAvailableCPUs)
    scores = instPool.map(runForClusterNum, [(cluster_num, copy.deepcopy(data), trials_to_do) for cluster_num in range(cluster_num_mi
    instPool.close()
    instPool.join()
```

**tagLowValuesDataframe**(df, cutoff, replacement)

```
Tag low values with replacement value.

Args:
    df: pandas DataFrame
    cutoff: values below the "cutoff" are replaced with "replacement" value
    replacement: replacement value

Returns:
    Processed pandas Dataframe

Usage:
    df_data = tagLowValuesDataframe(df_data, 1., 1.)
```

**tagMissingValuesDataframe**(df)

```
Tag missing (i.e. zero) values with NaN.

Args:
    df: pandas DataFrame

Returns:
    Processed pandas Dataframe

Usage:
    df_data = tagMissingValuesDataframe(df_data)
```

**timeSeriesClassification**(df_data, dataName, saveDir, hdf5fileName=None, p_cutoff=0.05, NumberOfRandomSamples=100000, NumberOfCPUs=4, frequencyBasedCl
calculateAutocorrelations=False, calculatePeriodograms=False)

```
Time series classification.

Args:
    df_data: pandas DataFrame
    dataName: data name, e.g. "myData_1"
    saveDir: path of directories poining to data storage
    hdf5fileName: preferred hdf5 file name and location
    p_cutoff: significance cutoff signals selection
    NumberOfRandomSamples: size of the bootstrap distribution to generate
    NumberOfCPUs: number of processes allowed to use in calculations
    frequencyBasedClassification: whether Autocorrelation of Frequency based
    calculateAutocorrelations: whether to recalculate Autocorrelations
    calculatePeriodograms: whether to recalculate Periodograms

Returns:
    None

Usage:
    timeSeriesClassification(df_data, dataName, saveDir, NumberOfRandomSamples = 10**5, NumberOfCPUs = 4, p_cutoff = 0.05, frequencyBa
```

**visualizeTimeSeriesClassification**(dataName, saveDir, numberOfLagsToDraw=3, hdf5fileName=None, exportClusteringObjects=False, writeClusteringObjectToBinari
AutocorrNotPeriodogr=True, vectorImage=True)

```
Visualize time series classification.

Args:
    dataName: data name
    saveDir: path of directories poining to data storage
    numberOfLagsToDraw: first top-N lags (or frequencies) to draw
    hdf5fileName: HDF5 storage path and name
    exportClusteringObjects: export clustering objects to xlsx files
    writeClusteringObjectToBinaries: export clustering objects to binary (pickle) files
    AutocorrNotPeriodogr: label to print on the plots

Returns:
```

### 15 PYIOMICA FUNCTION REFERENCES

```
        None

    Usage:
        visualizeTimeSeriesClassification('myData_1', '/dir1/dir2/', AutocorrNotPeriodogr=True, writeClusteringObjectToBinaries=True)
```

**write**(data, fileName, withPKLZextension=True, hdf5fileName=None, jsonFormat=False)

```
    Write object into a file. Pandas and Numpy objects are recorded in HDF5 format
    when 'hdf5fileName' is provided otherwise pickled into a new file.

    Args:
        data: data object to write into a file
        fileName: path of directories ending with the file name
        withPKLZextension: add ".pklz" to a pickle file
        hdf5fileName: path of directories ending with the file name. If None then data is pickled.
        jsonFormat: save data into compressed json file

    Returns:
        None

    Usage:
        write(exampleDataFrame, '/dir1/exampleDataFrame', hdf5fileName='/dir2/data.h5')
```

#### Data

**ConstantGeneDictionary** = None
**ConstantPyIOmicaDataDirectory** = '/Users/user/anaconda3/envs/pyiomicaTest/lib/python3.7/site-packages/pyiomica/data'
**ConstantPyIOmicaExampleVideosDirectory** = '/Users/user/anaconda3/envs/pyiomicaTest/lib/python3.7/site-packages/pyiomica/data/ExampleVideos'
**ConstantPyIOmicaExamplesDirectory** = '/Users/user/anaconda3/envs/pyiomicaTest/lib/python3.7/site-packages/pyiomica/data/ExampleData'
**PackageDirectory** = '/Users/user/anaconda3/envs/pyiomicaTest/lib/python3.7/site-packages/pyiomica'
**UserDataDirectory** = '/Users/user/anaconda3/envs/pyiomicaTest/lib/python3.7/site-packages/pyiomica/data'
**path** = '/Users/user/anaconda3/envs/pyiomicaTest/lib/python3.7/site-packages/pyiomica/data/ExampleVideos'
**readIn** = PosixPath('/Users/user/anaconda3/envs/pyiomica...thon3.7/site-packages/pyiomica/data/__init__.py')
